# Application of Supervised Machine Learning Models for Drug-Action Prediction Towards Nuclear Type I Receptors

**DOI:** 10.1101/2024.05.03.592421

**Authors:** Rajeev Jaundoo, Jack A. Tuszynski, Travis J.A. Craddock

**Affiliations:** University of Alberta; Nova Southeastern University

## Abstract

Interactions between drugs can lead to adverse side effects for patients taking combination therapies to treat complex diseases such as cancer. Knowledge of drug-action towards a receptor would allow these drug-drug interactions to be predicted, and in this study, we trained a total of 5 different machine learning models to classify whether a given drug was an agonist (activator), antagonist (blocker), or a decoy (non-binder) to each of the androgen, estrogen, glucocorticoid, and progesterone receptors. The classification performance and efficiency, measured in training time, of the decision tree, naïve Bayes, neural network, random forest, and support vector machine models for each receptor were then compared. The results showed that the decision tree and naïve Bayes models were best suited for drug-action prediction across all receptors while only requiring minutes of training time at most. Future work will focus on increasing the prediction accuracy of antagonist drugs, integrating experimental data during training, and using other targets outside of nuclear type I receptors.

## 2. Introduction

Complex diseases such as cancer involve numerous pathways and receptors, which leads to patients taking drug combination therapies to combat the symptoms of their disease. Unfortunately, drugs can alter the absorption, distribution, metabolism, and/or excretion of one another, which may result in adverse drug-drug interactions (Becker, 2011). Drug-action prediction, that is, classifying a drug either an agonist, antagonist, or non-binder/decoy to a target, would help mitigate this risk, and additionally, would be useful for drug repurposing, a process in which existing pharmaceuticals are used to treat a disease or indication not originally intended. Furthermore, drug libraries such as ZINC (Irwin et al., 2020) that contain millions of compounds could easily be filtered for further analysis (e.g., drug docking) based on drug-action towards the target(s) of interest.

Machine learning (ML) models have gained a significant amount of popularity in recent years for completing tasks such as predicting the binding affinity and bound conformation of a ligand to a target (Yang et al., 2022; Isert et al., 2023), rapid screening of drug libraries based on specific properties including Lipinski’s rule-of-5 (Cáceres et al., 2020), and predicting the 3-D structures of proteins from only their amino acid sequences (e.g., AlphaFold) among others. In this study, we focused on developing ML models to predict drug-action towards a set of targets. An agonist induces a biological response upon binding to a receptor, while antagonists reduce or block the action of another drug, often agonists, by occupying the same binding region on the receptor (Neubig et al., 2003).

Only full agonists were considered while partial and irreversible agonists were not, and for antagonists, both competitive and non-competitive types were used. The targets were defined as the androgen (AR), estrogen (ER), glucocorticoid (GR), and progesterone (PR) nuclear type I receptors, while the decision tree (DECTRE), naïve Bayes (NAIBAY), neural network (NEUNET), random forest (RANFOR), and support vector machine (SVM) learners were trained to compare their performance as well as efficiency, or the total time required for training.

## 3. Methods

As an overview, published literature was first used to find experimental agonists and antagonists of AR, ER, GR, and PR, while decoys of all receptors were obtained from the Database of Useful Decoys: Enhanced (DUD-E; Mysinger et al., 2012). Next, the agonists, antagonists, and decoys for each receptor were split 50:50 into training and test sets, and in cases with odd numbers, the extra drug was placed into the training set. A process known as data augmentation was then performed to increase the total number of agonists and antagonists for each receptor. Afterwards, the Molecular Operating Environment (MOE; Chemical Computing Group, 2019a) application was employed to generate features of all drugs, including the total number of carbon, nitrogen, sulfur, etc. atoms, Lipinski’s Rule-of-5 violations, and torsion energy among many others. ML was completed using RapidMiner Studio (version 9.6; Mierswa & Klinkenberg, 2020), where cross-validation (CV) was utilized during the training process. Finally, the performance of each model was assessed using the validation set, where the precision and recall metrics were used to quantify each model’s accuracy in classifying drugs as either agonists, antagonists, or decoys. These metrics accounted for the number of true positives (TPs), false positives (FPs), and false negatives (FNs) within each model’s predictions. See Equations (1 and (2 below.

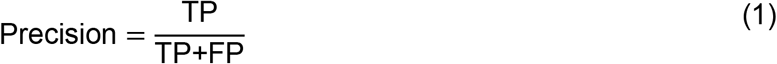

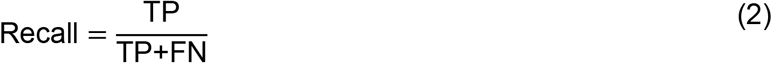

**Figure 1:**
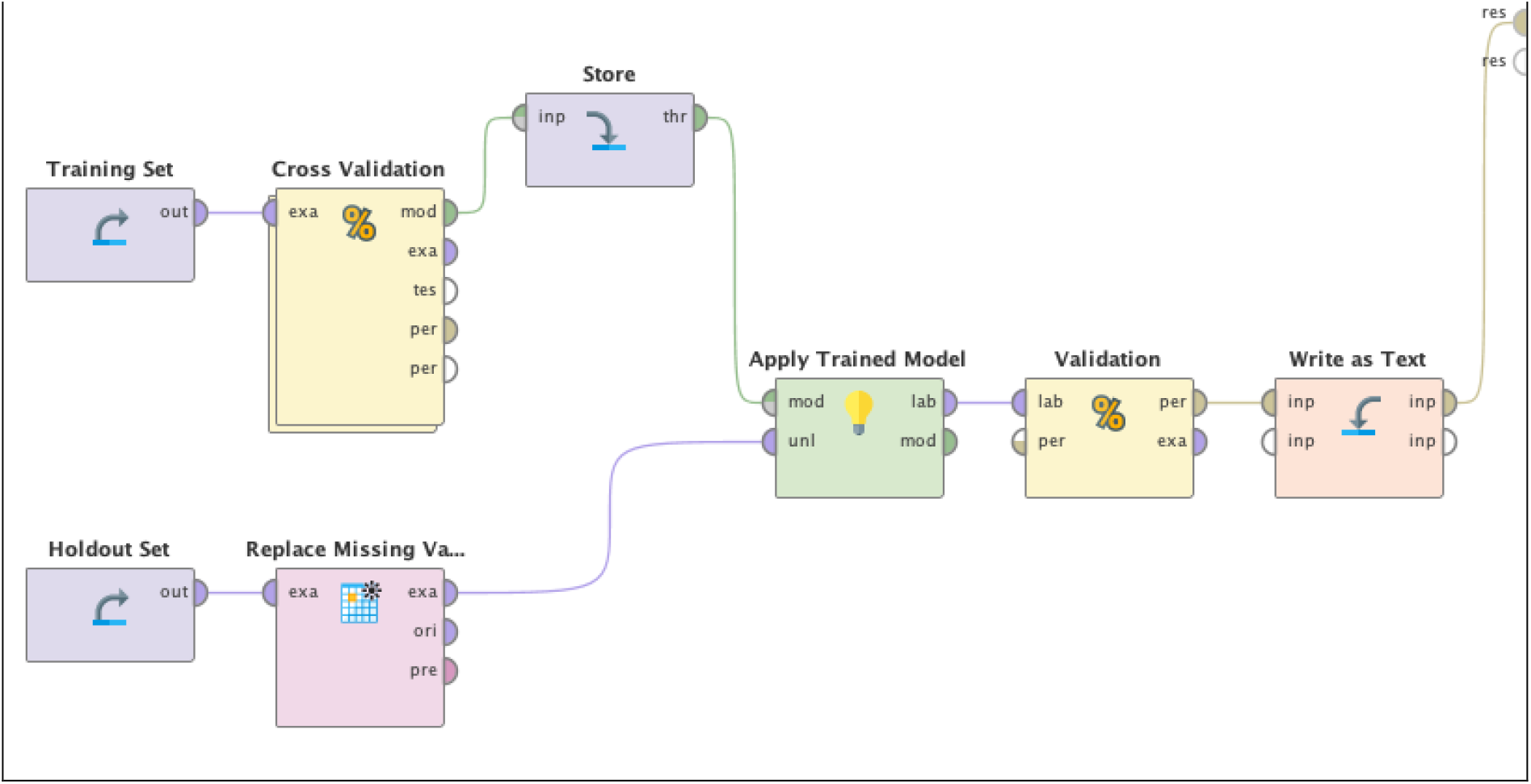
Overview of the ML process for each learner. A model was generated using the “Cross Validation” operator that utilized the entire training set, while CV was completed to assess its performance. The model was then tested using the validation dataset to calculate its final performance. Note the “Replace Missing Va…[lues]” operator that replaced all missing entries within both the training (not pictured) and test set with −1.

### 3.1. Agonists, Antagonists, & Decoys

Published literature as well as expert curated data sources such as the Protein Data Bank (PDB), DrugBank, ChEMBL, and ChemSpider were explored to obtain agonist and antagonist drugs for AR, ER, GR, and PR. Here, the simplified molecular-input line entry system (SMILES) string was downloaded for each drug, which represents its atoms and stereochemistry (O’Boyle, 2012). However, there were relatively few agonists and antagonists, often less than 100, compared to the total number of decoys available (>14,000) for each receptor; see Table 1. This could have led to overfitting, which occurs when the training data provides a poor representation of the actual population of agonists and/or antagonists, leading to low performance on novel data (Mutasa et al., 2020).

**Table 1:** Total number of agonists, antagonists, and decoys obtained for each receptor.

| Receptor | Agonists | Antagonists | Decoys |
| --- | --- | --- | --- |
| AR | 106 | 18 | 14,503 |
| ER | 62 | 28 | 20,818 |
| GR | 44 | 10 | 15,185 |
| PR | 45 | 7 | 15,814 |

**Table 2:** Total number of agonists within the training and test data sets before and after data augmentation.

|  | Training Set |  | Test Set |  |
| --- | --- | --- | --- | --- |
|  | *Before* | *After* | *Before* | *After* |
| AR | 53 | 2198 | 53 | 1579 |
| ER | 31 | 1440 | 31 | 1417 |
| GR | 22 | 4014 | 22 | 3437 |
| PR | 23 | 1434 | 22 | 799 |

To address this issue, data augmentation was performed to increase the number of agonists and antagonists, which involved generating numerous conformations of both sets. While 2-D properties of these generated conformations such as number of carbon atoms or number of aromatic rings would be the same as the original, 3-D properties including the torsion and bond stretch energy would be different, increasing the variation within the population of agonists and antagonists of the training set. This process was completed using the “Conformation Input” tool in MOE.

Conformations were generated in 5 main steps. First, acids or bases that were previously (de)protonated were corrected. Each drug was then broken into overlapping fragments, where the conformation of each was determined using a stochastic conformational search (Chemical Computing Group, 2019b). These fragments were then rebuilt to create numerous conformations of the same agonist/antagonist, and the van der Waals (vdW) energy of each was calculated to filter those with bad contacts between atoms. Finally, the strain energies were calculated and, along with the conformations of each agonist and antagonist, were written to the output file (Chemical Computing Group, 2019b).

**Table 3:** Total number of antagonists within the training and test data sets before and after data augmentation.

|  | Training Set |  | Test Set |  |
| --- | --- | --- | --- | --- |
|  | *Before* | *After* | *Before* | *After* |
| AR | 9 | 588 | 9 | 853 |
| ER | 14 | 2370 | 14 | 1337 |
| GR | 5 | 542 | 5 | 477 |
| PR | 4 | 584 | 3 | 258 |

### 3.2. Features

Features are properties or characteristics that describe a given agonist, antagonist, or decoy such as its molecular weight for example, and are used during training to differentiate between the classes. A total of 435 features were generated by MOE via the QuaSAR-Descriptor tool, which included 2-D descriptors (e.g., number of aromatic rings and heavy, H-bond donor, etc. atoms; number of single rotatable, triple, etc. bonds), internal 3-D descriptors (e.g., total, potential, electrostatic, etc. energy), external 3-D descriptors (e.g. dipole moment, non-bonded interaction energy, etc.), and protein descriptors (e.g., net charge, hydrophobicity, volume, etc.). Supplementary Table 1 contains a full list of all descriptors.

### 3.3. Machine Learning

Training of the DECTRE, NAIBAY, NEUNET, RANFOR, and SVM learners was completed using *k*-fold CV, a resampling method that partitions the entire training data into *k* subsets, where *k* = 10 in this study. Here, *k* − 1 subsets were used for training while the last subset was kept for validation. This process is repeated *k* number of times (folds) such that each subset is used as a validation set exactly once. Stratified sampling was used as well, meaning that both the training and validation sets within each fold of CV contained approximately the same proportion of agonists, antagonists, and decoys as the entire training data. Missing values were replaced with −1.

Moreover, in each fold of CV, feature selection was first performed before training to keep only the most relevant features. This process reduces noise and can increase performance by allowing the learner to distinguish relationships more easily between each class and the remaining features (Hall & Smith, 1998). Here, the forward algorithm was employed for feature selection. The final feature set was first populated by evaluating each feature individually using CV to identify the one with the best performance. Next, the remaining features were individually coupled with the existing set to identify the one that led to the greatest performance increase. This process was repeated until there was either no increase in performance or all features were evaluated.

To summarize the overall process, training was performed using 10-fold CV, where the training subsets of each fold were used to complete feature selection before training and validation. After all 10 folds were finished, the average performance of each fold’s model was calculated. Note that the final output model was based on the entire training dataset, where its performance was assessed using the validation set.

## 4. Results

The performance of all models was calculated using the accuracy metric, ^Correct^/Total, which is the total number of correctly predicted agonists, antagonists, and decoys divided by the total number of drugs for that receptor. Additionally, the baseline for each receptor was calculated by dividing the number of entries within the largest class, in this case decoys, and dividing it by the total number of agonists, antagonists, and decoys available for each of AR, ER, GR, and PR. The performance of all trained models needed to be higher than baseline, otherwise their predictions would have been no different than simply classifying all drugs into the largest class. See Table 4.

**Table 4:** Baseline percentages for each receptor.

| *Receptor* | *Baseline Percent* |
| --- | --- |
| AR | 75% |
| ER | 79% |
| GR | 66% |
| PR | 88% |

### 4.1. Performance: AR

All models had similar performance in classifying decoys compared to the actual or ground truth (TRUE) number, although SVM had more FNs for agonists and antagonists. All models had very high recall for decoys, the lowest being 99.96% (NAIBAY) and the highest being 100% (DECTRE), while the precision for this class ranged from 77.17% (SVM) to 97.24% (NAIBAY). For antagonists, SVM had the worst precision and recall at 0% while NAIBAY had the best at 56.51% and 87.64% respectively. Finally, NEUNET was able to best classify agonists with a recall of 94.49%, while SVM was shown to have the best precision at 99.65%. The model with the lowest recall for agonists was RANFOR at 78.02% and NAIBAY had the lowest precision at 85.41%. See Supplementary Tables 2-6 for the precision and recall of each model.

**Figure 2.**
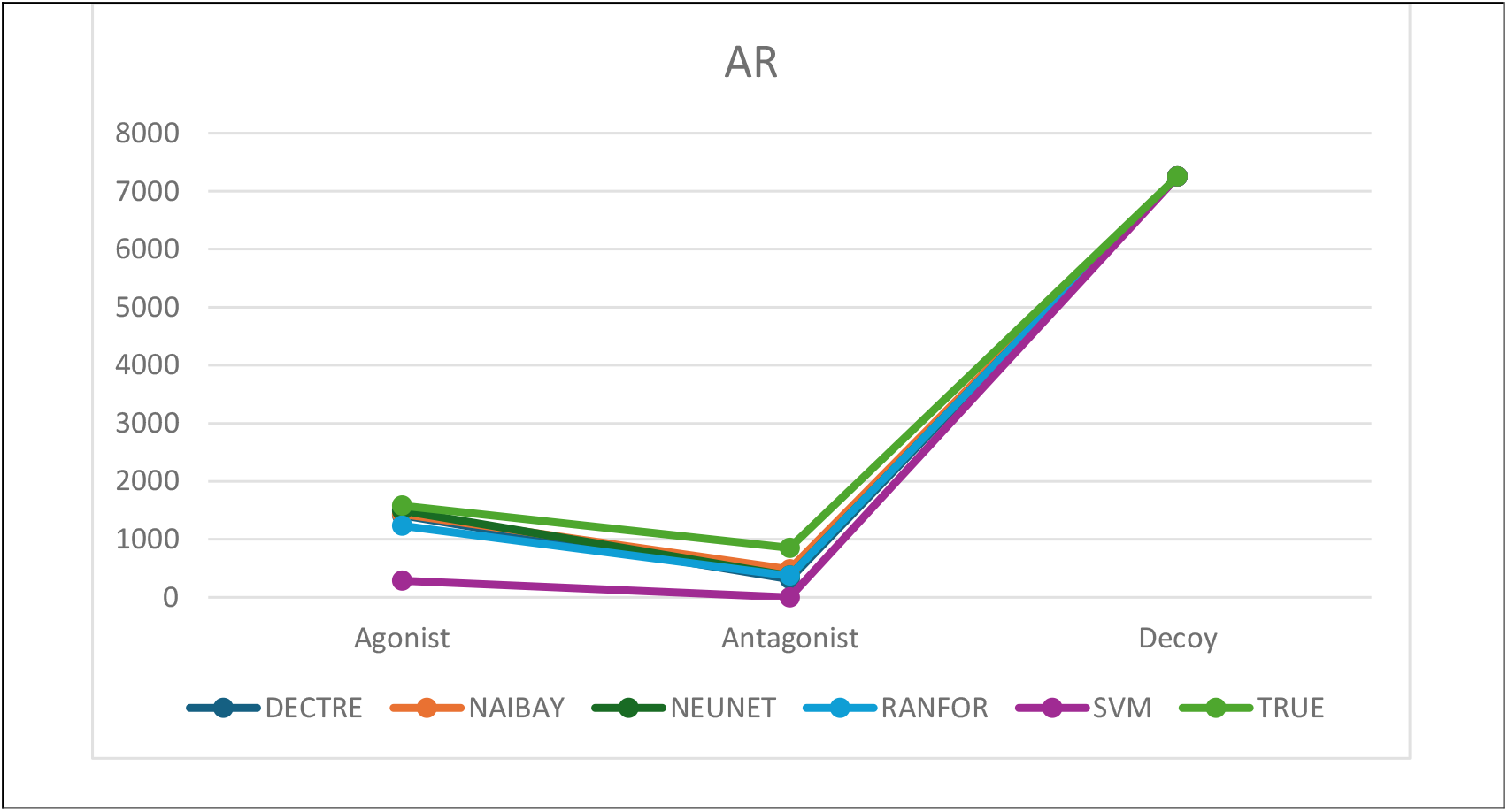
The performance of each model compared to the ground truth (TRUE) in classifying agonists, antagonists, and decoys for AR.

### 4.2. Performance: ER

All models except for DECTRE struggled to classify agonists as shown by their recall that ranged from 4.09% (SVM) to 90.54%. That being said, DECTRE had the worst agonist precision at 75.96% while NEUNET had the best at 96.02% followed by SVM (92.06%). Both NAIBAY and NEUNET were shown to be close to TRUE regarding antagonist recall, 87.96% and 90.73% respectively, while the other models ranged from 18.70% (SVM) to 37.70% (DECTRE). SVM performed especially well in antagonist class precision at 98.04%, seeing as the second highest was DECTRE at 80% and the worst was RANFOR (25.25%). For decoys, all models performed similarly, the lowest recall was 99.95% for DECTRE while the best percentage was from the SVM model at 99.97%. Precision for the decoy class had a more variable range with a low of 81.01% for SVM to a high of 99.20% for NEUNET. See Supplementary Tables 7-11.

**Figure 3.**
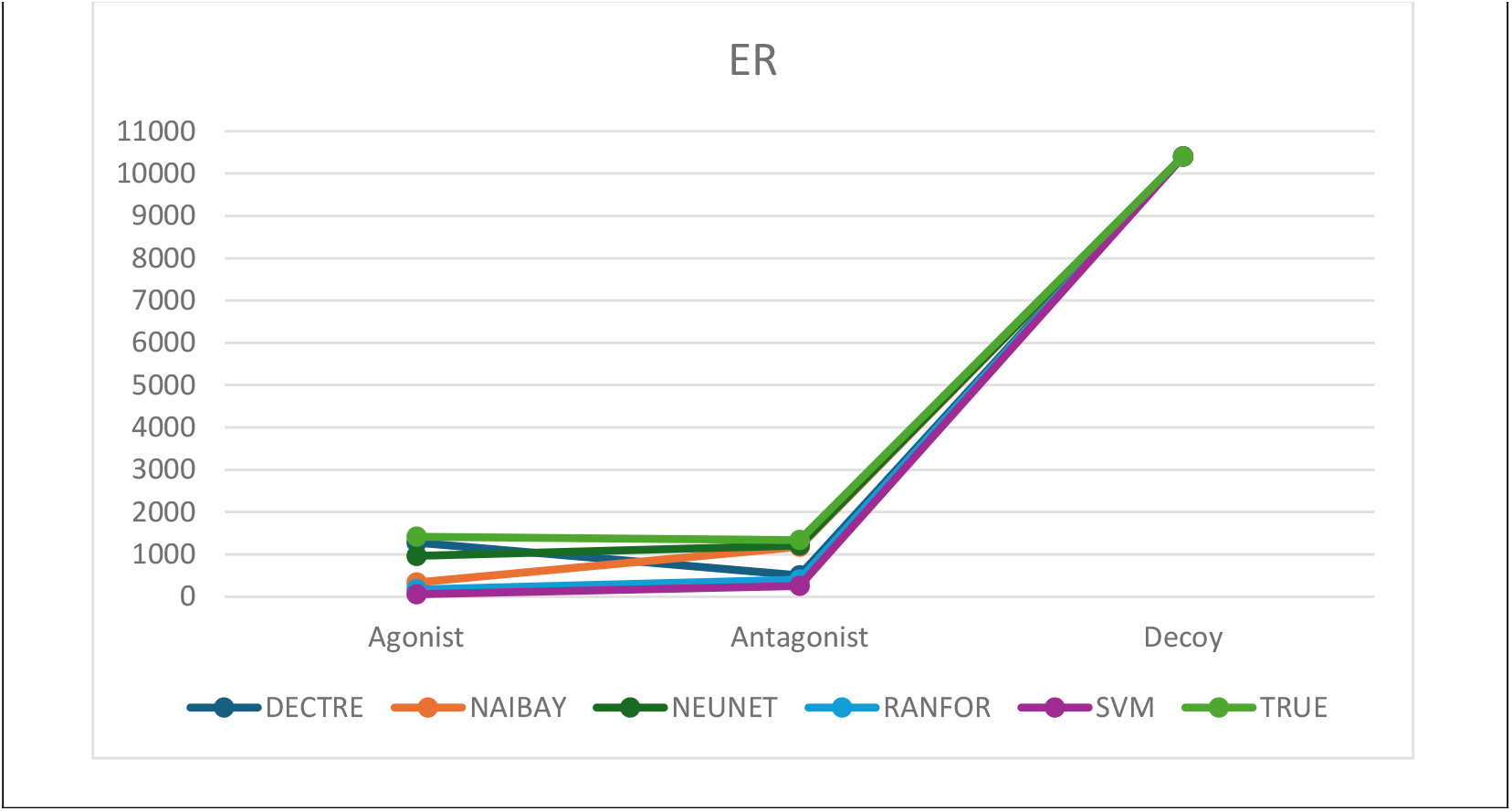
The performance of each model compared to the ground truth (TRUE) in classifying agonists, antagonists, and decoys for ER.

### 4.3. Performance: GR

DECTRE, NAIBAY, and NEUNET correctly classified all agonists with 100% recall. Agonist precision had a low of 92.37% for the NAIBAY model with a range of 99.37% (RANFOR) to 99.83% (NEUNET) on the high end. All models performed poorly in antagonist recall, the worst was SVM at 0% while DECTRE, NAIBAY, and RANFOR were 5.45%, and NEUNET ranked best at 12.37%. The precision for the antagonist class was 100% for DECTRE, NEUNET, and RANFOR, followed by NAIBAY at 81.25%, and SVM at 0%. Both recall and precision for the decoy class fared better for all models, with a recall ranging from 99.91% at the lowest (NAIBAY) and 100% at best (DECTRE). SVM had the worst precision for this class at 80.60% and NAIBAY had the highest: 97.83%. Supplementary Tables 12-16 display the performance of all models for GR.

**Figure 4.**
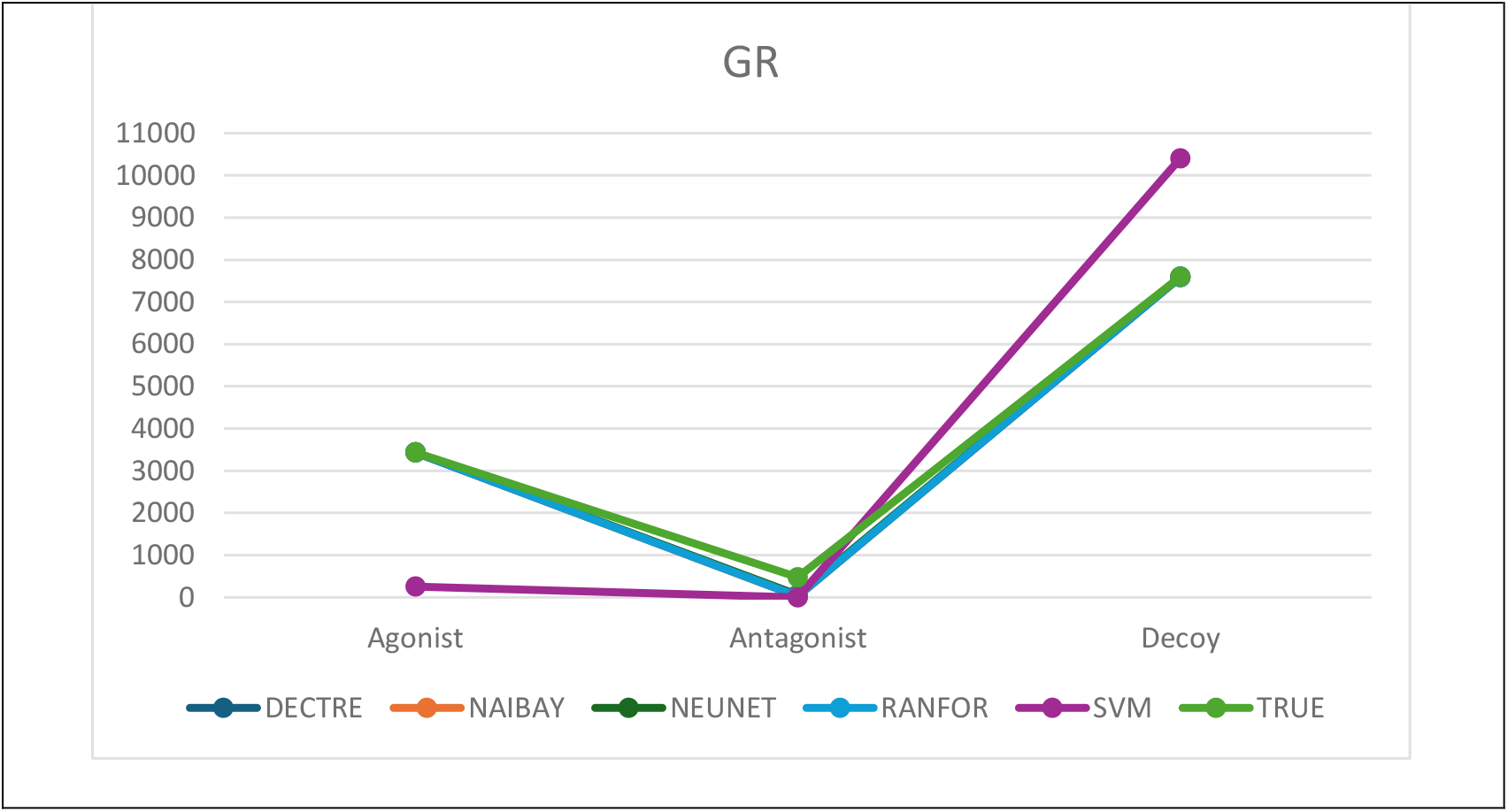
The performance of each model compared to the ground truth (TRUE) in classifying agonists, antagonists, and decoys for GR.

### 4.4. Performance: PR

All models, with the exception of SVM, were mainly consistent with TRUE in classifying agonists, antagonists, and decoys. The agonist class had a recall of 98.75% for NEUNET and RANFOR, 99.25% for DECTRE, and 99.50% for the NAIBAY model. SVM was the worst at 23.03% but had a precision of 100% for the agonist class, same as DECTRE. The lowest agonist class precision was from the NEUNET model at 87.57% while NAIBAY and RANFOR were 99.62% and 99.87% respectively. DECTRE, NAIBAY, and RANFOR models all had 100% recall for antagonists, followed by NEUNET (56.59%) and then SVM in last at 38.87%. Similarly, SVM (67.57%) and NEUNET (96.05%) were also ranked last in antagonist precision, with NAIBAY being first at 99.61%. The recall for decoys was within 99.95% to 100% for all models, where NAIBAY ranked last, and the lowest decoy precision was SVM at 91.58%, while NAIBAY, NEUNET, and RANFOR were 99.95% and DECTRE was 100%. See Supplementary Tables 17-21 for the precision and recall of all trained models.

**Figure 5.**
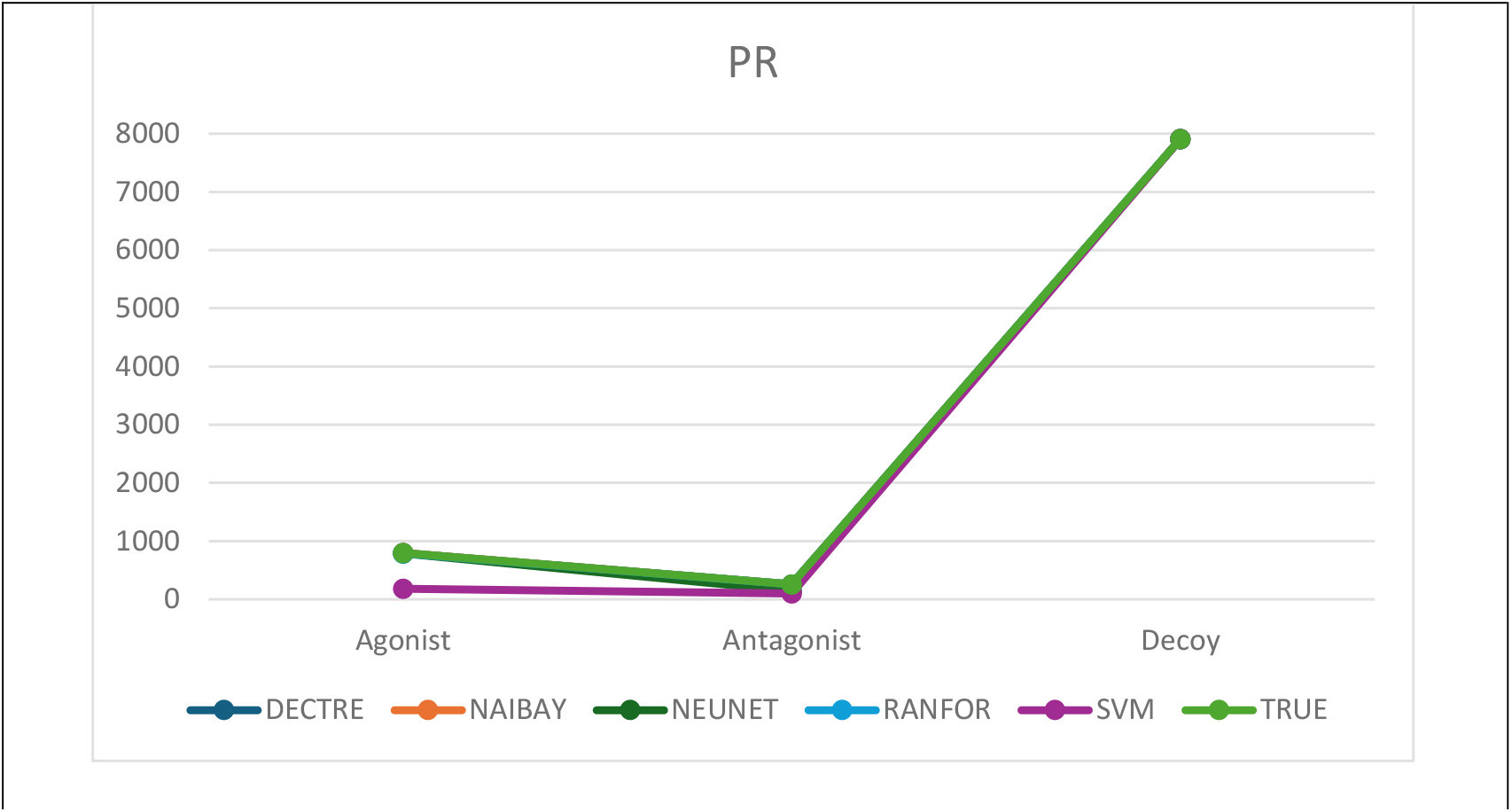
The performance of each model compared to the ground truth (TRUE) in classifying agonists, antagonists, and decoys for PR.

### 4.5. Summary

To summarize, the model with the best performance for each receptor was: NAIBAY for AR, NEUNET for ER and GR, and finally, DECTRE and NAIBAY were within 0.01% of each other for PR. See Table 5 below.

**Table 5:** Performance of all learners on all receptors.

|  | DECTRE | NAIBAY | NEUNET | RANFOR | SVM |
| --- | --- | --- | --- | --- | --- |
| AR | 92.73% | 94.64% | 94.04% | 91.41% | 77.83% |
| ER | 92.61% | 90.51% | 95.59% | 83.43% | 81.39% |
| GR | 96.08% | 96.02% | 96.32% | 96.03% | 80.93% |
| PR | 99.92% | 99.91% | 98.64% | 99.87% | 91.36% |
| Average | 95.34% | 95.27% | 96.16% | 92.69% | 82.88% |

**Table 6:**
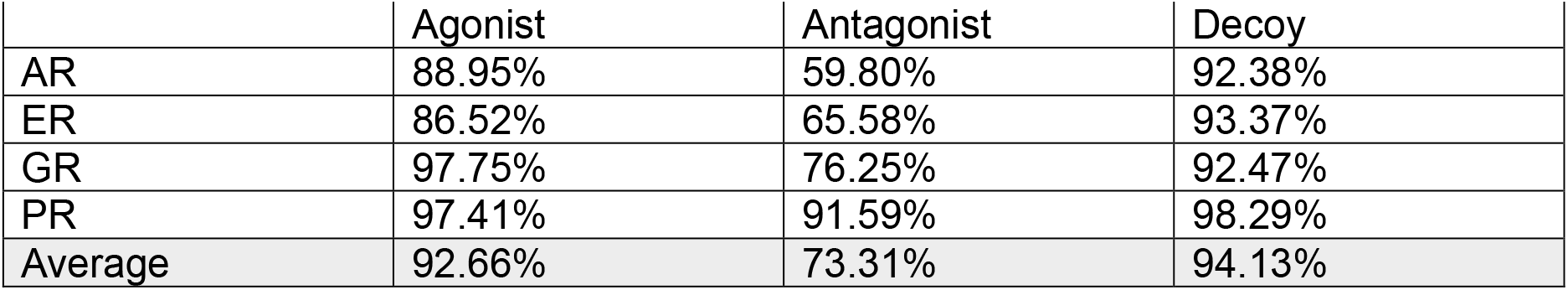
Average class precision of the agonist, antagonist, and decoy classes for all learners on all receptors.

|  | Agonist | Antagonist | Decoy |
| --- | --- | --- | --- |
| AR | 88.95% | 59.80% | 92.38% |
| ER | 86.52% | 65.58% | 93.37% |
| GR | 97.75% | 76.25% | 92.47% |
| PR | 97.41% | 91.59% | 98.29% |
| Average | 92.66% | 73.31% | 94.13% |

In regard to training time, Table 8 shows NAIBAY had the fastest average over all receptors with DECTRE trailing closely behind. The slowest learner was by far SVM, which happened to have the worst performance as well.

**Table 7:** Average class recall of the agonist, antagonist, and decoy classes for all learners on all receptors.

|  | Agonist | Antagonist | Decoy |
| --- | --- | --- | --- |
| AR | 74.21% | 36.59% | 99.98% |
| ER | 39.70% | 53.09% | 99.95% |
| GR | 83.56% | 05.74% | 99.94% |
| PR | 83.86% | 79.07% | 99.98% |
| Average | 70.33% | 43.62% | 99.96% |

**Table 8:** Total run-time of all learners on all receptors. The format used is day:hour:minute:second.

|  | DECTRE | NAIBAY | NEUNET | RANFOR | SVM |
| --- | --- | --- | --- | --- | --- |
| AR | 00:00:03:33 | 00:00:01:09 | 00:06:34:47 | 00:02:21:18 | 01:16:47:47 |
| ER | 00:00:13:39 | 00:00:02:25 | 00:21:38:12 | 00:08:01:19 | 15:12:40:35 |
| GR | 00:00:01:35 | 00:00:01:06 | 00:08:46:01 | 00:04:20:41 | 13:22:40:27 |
| PR | 00:00:02:58 | 00:00:01:00 | 00:04:14:59 | 00:01:19:57 | 06:02:55:33 |
| Average | 00:00:05:27 | 00:00:01:25 | 00:10:18:30 | 00:04:03:49 | 09:07:46:06 |

## 5. Discussion

Overall, the NEUNET model had the best average performance of the 5 models, followed by RANFOR, NAIBAY, DECTRE, and then SVM; see Table 5. A 2-tailed paired samples t-test was used to determine if the predicted classes between 2 different models were significantly different to one another if *p* < 0.05. The performance of NEUNET over AR, ER, GR, and PR was found to not be statistically significant compared to RANFOR (p =0.33), NAIBAY (p =0.58), or DECTRE (p = 0.43). Additionally, the performance between the DECTRE and NAIBAY (p=0.94), DECTRE and RANFOR (p=0.31), and NAIBAY and RANFOR (p=0.22) models were not significant either. Only SVM was found to be significant compared to the other models: DECTRE (p=0.00), NAIBAY (p=0.01), NEUNET (p=0.01), and RANFOR (p=0.04), which is consistent with its generally poor precision and recall compared to these other models.

Training time also varied between all learners, see Table 8 and Figure 6. Here, DECTRE and NAIBAY took minutes across all receptors while NEUNET, which performed approximately 1% better than both, took multiple hours in comparison. Moreover, SVM had the longest training time, measured in days rather than hours, and ER required the most training time for all learners compared to the other AR, GR, and PR targets. Regarding precision, which measured the number of drugs that were correctly classified as an agonist, antagonist, or decoy, the decoy class had the best/highest percentage, followed by agonists and then antagonists; see Table 6. Similarly, Table 7 shows that antagonists also had the worst averaged recall, which is the number of drugs correctly identified as agonists, antagonists, or decoys while accounting for FNs, while the decoy class had the best percentage and agonists were again ranked in the middle.

**Figure 6.**
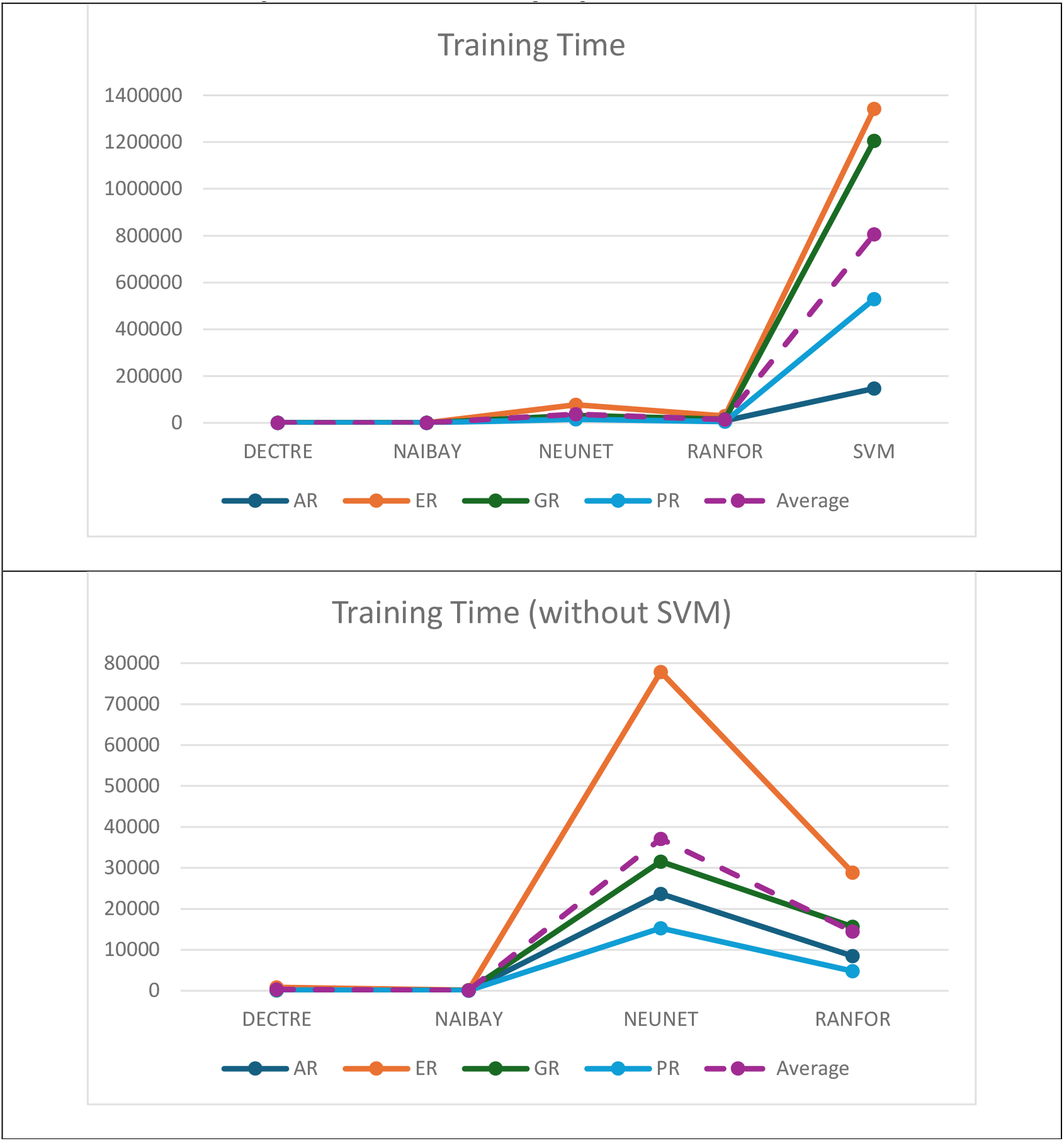
The total training/run-time (in seconds) of the models, with and without the inclusion of SVM, over all receptors. The average time of all models on all receptors are shown in purple with dashed lines.

These results may have been due to the fact the antagonist class for AR, GR, and PR had fewer entries compared to the agonist and decoy classes, the latter of which contained the most examples for training and validation across all receptors. Furthermore, differences between agonists and antagonists may be subtle or inconsistent, making identification of antagonists that much harder. Interestingly, the comparison of learners in Table 5 shows that DECTRE performed better than RANFOR over all receptors. RANFOR is a meta-learner comprised of multiple DECTREs, where the majority consensus determines the final predicted class. Bootstrap sampling may have been the cause of this discrepancy, where the data used to generate each DECTRE in RANFOR were not representative of the actual population; for example, some DECTREs may have been trained using mostly agonist/antagonist/decoy data instead of evenly distributed datasets. However, the biggest limitation of this study was the required use of data augmentation due to the few agonists and antagonists found in literature. That being said, even if the differences in 3-D features were not enough to differentiate between the original agonist/antagonist and their conformations, no cross-contamination would have occurred between the training and validation sets because the split between these datasets occurred prior to data augmentation.

## 6. Conclusion

In this study, the DECTRE, NAIBAY, NEUNET, RANFOR, and SVM learners were trained to identify whether a given drug would be an agonist, antagonist, or decoy to each of the AR, ER, GR, and PR targets. The performance and required training time of each model on every receptor was compared to determine the one with a balance of both accuracy and efficiency. Drug-action prediction can be used for a variety of tasks including filtering of drug libraries, lead optimization, and prediction of drug interactions to name a few.

The results showed all models were able to exceed the baseline performance (Table 4) for every receptor, or in other words, all models were more accurate than simply classifying all query drugs into the largest class. Moreover, there was no statistical difference between any of the top 3 models: NEUNET and DECTRE (*p* = 0.43), NEUNET and NAIBAY (*p* = 0.58), and NAIBAY and DECTRE (*p* = 0.94), meaning these models had no significant difference to one another in classification of agonist, antagonist, and decoy drugs. Keeping this in mind, the training time for both DECTRE and NAIBAY were measured in minutes while NEUNET required hours for less than a 1% increase in averaged performance.

Future work will focus on the improvement of antagonist classification, which was poor compared to the agonist and decoy classes. This will be accomplished by increasing the total number of antagonists for each receptor as well as generating more features to allow the learners to better differentiate between the classes. The paradigm shift from conventional *in-silico* tools to ML allows researchers to take advantage of existing biomedical data to generate models that require less resources to run compared to drug docking or molecular dynamics simulations for instance. The methodology used in this study can now be extended to various other targets, furthering efforts in drug development and repurposing in order to treat diseases.

## Supporting information

All Supplementary Tables

## References

Becker, D. E. (2011). Adverse drug interactions. Anesthesia progress, 58(1), 31–41. 10.2344/0003-3006-58.1.31

Cáceres, E. L., Tudor, M., & Cheng, A. C. (2020). Deep learning approaches in predicting ADMET properties. Future Medicinal Chemistry, 12(22), 1995–1999. 10.4155/fmc-2020-0259

Chemical Computing Group. (2019a). Molecular Operating Environment (MOE). 2019.01.

Chemical Computing Group. (2019b). MOE User Guide.

Hall, M. A., & Smith, L. A. (1998). Practical feature subset selection for machine learning (C. McDonald, Ed.; pp. 181–191). Springer.

Irwin, J. J., Tang, K. G., Young, J., Dandarchuluun, C., Wong, B. R., Khurelbaatar, M., Moroz, Y.S., Mayfield, J., & Sayle, R. A. (2020). ZINC20—a free ultralarge-scale chemical database for ligand discovery. Journal of chemical information and modeling, 60(12), 6065–6073. 10.1021/acs.jcim.0c00675

Isert, C., Atz, K., & Schneider, G. (2023). Structure-based drug design with geometric deep learning. Current Opinion in Structural Biology, 79, 102548. 10.1016/j.sbi.2023.102548

Mierswa, I., & Klinkenberg, R. (2020). RapidMiner Studio (version 9.6).

Mutasa, S., Sun, S., & Ha, R. (2020). Understanding artificial intelligence based radiology studies: What is overfitting? Clinical Imaging, 65, 96–99. 10.1016/j.clinimag.2020.04.025

Mysinger, M. M., Carchia, M., Irwin, J. J., & Shoichet, B. K. (2012). Directory of useful decoys, enhanced (DUD-E): better ligands and decoys for better benchmarking. Journal of medicinal chemistry, 55(14), 6582–6594. 10.1021/jm300687e

Neubig, R. R., Spedding, M., Kenakin, T., & Christopoulos, A. (2003). International Union of Pharmacology Committee on Receptor Nomenclature and Drug Classification. XXXVIII. Update on terms and symbols in quantitative pharmacology. Pharmacological Reviews, 55(4), 597–606. 10.1124/pr.55.4.4

O’Boyle, N. M. (2012). Towards a Universal SMILES representation - A standard method to generate canonical SMILES based on the InChI. Journal of Cheminformatics, 4(1), 1–14. 10.1186/1758-2946-4-22

Yang, C., Chen, E. A., & Zhang, Y. (2022). Protein–Ligand docking in the machine-learning era. Molecules, 27(14), 4568. 10.3390/molecules27144568

