## Supplementary material for "Application of Supervised Machine Learning Models for Drug-Action Prediction Towards Nuclear Type I Receptors": All Supplementary Tables

| **Supplementary Table 1: All 435 features generated from MOE. The code represents the name of the feature, the class denotes whether the given feature is a 2D, internal 3d (i3D) or external 3d (x3D) type.** | | |
| --- | --- | --- |
| *Code* | *Class* | *Description* |
| AM1_dipole | i3D | Dipole moment |
| AM1_E | i3D | Total energy (kcal/mol) |
| AM1_Eele | i3D | Electronic energy (kcal/mol) |
| AM1_HF | i3D | Heat of formation (kcal) |
| AM1_HOMO | i3D | HOMO energy (eV) |
| AM1_IP | i3D | Ionization potential (eV) |
| AM1_LUMO | i3D | LUMO energy (eV) |
| apol | 2D | Sum of atomic polarizabilities |
| ASA | i3D | Water accessible surface area |
| ASA+ | i3D | Positive accessible surface area |
| ASA- | i3D | Negative accessible surface area |
| ASA_H | i3D | Total hydrophobic surface area |
| ASA_P | i3D | Total polar surface area |
| ast_fraglike | 2D | Astex Fragment-like Test |
| ast_fraglike_ext | 2D | Astex Fragment-like Test (Extended) |
| ast_violation | 2D | Astex Fragment-like Violation Count |
| ast_violation_ext | 2D | Astex Fragment-like Violation Count (Extended) |
| a_acc | 2D | Number of H-bond acceptor atoms |
| a_acid | 2D | Number of acidic atoms |
| a_aro | 2D | Number of aromatic atoms |
| a_base | 2D | Number of basic atoms |
| a_count | 2D | Number of atoms |
| a_don | 2D | Number of H-bond donor atoms |
| a_donacc | 2D | Number of H-bond donor + acceptor atoms |
| a_heavy | 2D | Number of heavy atoms |
| a_hyd | 2D | Number of hydrophobic atoms |
| a_IC | 2D | Atom information content (total) |
| a_ICM | 2D | Atom information content (mean) |
| a_nB | 2D | Number of boron atoms |
| a_nBr | 2D | Number of bromine atoms |
| a_nC | 2D | Number of carbon atoms |
| a_nCl | 2D | Number of chlorine atoms |
| a_nF | 2D | Number of fluorine atoms |
| a_nH | 2D | Number of hydrogen atoms |
| a_nI | 2D | Number of iodine atoms |
| a_nN | 2D | Number of nitrogen atoms |
| a_nO | 2D | Number of oxygen atoms |
| a_nP | 2D | Number of phosphorus atoms |
| a_nS | 2D | Number of sulfur atoms |
| balabanJ | 2D | Balaban averaged distance sum connectivity |
| BCUT_PEOE_0 | 2D | PEOE Charge BCUT (0/3) |
| BCUT_PEOE_1 | 2D | PEOE Charge BCUT (1/3) |
| BCUT_PEOE_2 | 2D | PEOE Charge BCUT (2/3) |
| BCUT_PEOE_3 | 2D | PEOE Charge BCUT (3/3) |
| BCUT_SLOGP_0 | 2D | LogP BCUT (0/3) |
| BCUT_SLOGP_1 | 2D | LogP BCUT (1/3) |
| BCUT_SLOGP_2 | 2D | LogP BCUT (2/3) |
| BCUT_SLOGP_3 | 2D | LogP BCUT (3/3) |
| BCUT_SMR_0 | 2D | Molar Refractivity BCUT (0/3) |
| BCUT_SMR_1 | 2D | Molar Refractivity BCUT (1/3) |
| BCUT_SMR_2 | 2D | Molar Refractivity BCUT (2/3) |
| BCUT_SMR_3 | 2D | Molar Refractivity BCUT (3/3) |
| bpol | 2D | Difference of bonded atom polarizabilities |
| b_1rotN | 2D | Number of rotatable single bonds |
| b_1rotR | 2D | Fraction of rotatable single bonds |
| b_ar | 2D | Number of aromatic bonds |
| b_count | 2D | Number of bonds |
| b_double | 2D | Number of double bonds |
| b_heavy | 2D | Number of heavy-heavy bonds |
| b_max1len | 2D | Maximum single-bond chain length |
| b_rotN | 2D | Number of rotatable bonds |
| b_rotR | 2D | Fraction of rotatable bonds |
| b_single | 2D | Number of single bonds |
| b_triple | 2D | Number of triple bonds |
| CASA+ | i3D | Charge-weighted positive surface area |
| CASA- | i3D | Charge-weighted negative surface area |
| chi0 | 2D | Atomic connectivity index (order 0) |
| chi0v | 2D | Atomic valence connectivity index (order 0) |
| chi0v_C | 2D | Carbon valence connectivity index (order 0) |
| chi0_C | 2D | Carbon connectivity index (order 0) |
| chi1 | 2D | Atomic connectivity index (order 1) |
| chi1v | 2D | Atomic valence connectivity index (order 1) |
| chi1v_C | 2D | Carbon valence connectivity index (order 1) |
| chi1_C | 2D | Carbon connectivity index (order 1) |
| chiral | 2D | Number of chiral centers |
| chiral_u | 2D | Number of unconstrained chiral centers |
| DASA | i3D | Absolute difference in surface area |
| DCASA | i3D | Absolute difference in charge-weighted areas |
| dens | i3D | Mass density (AMU/A^3) |
| density | 2D | Mass density (AMU/A**3) |
| diameter | 2D | Largest vertex eccentricity in graph |
| dipole | i3D | Dipole moment |
| dipoleX | x3D | Dipole moment (X) |
| dipoleY | x3D | Dipole moment (Y) |
| dipoleZ | x3D | Dipole moment (Z) |
| E | i3D | Potential Energy |
| E_ang | i3D | Angle Bend Energy |
| E_ele | i3D | Electrostatic energy |
| E_nb | i3D | Non-bonded energy |
| E_oop | i3D | Out-of-plane Energy |
| E_rele | x3D | Electrostatic Interaction Energy |
| E_rnb | x3D | Non-bonded Interaction Energy |
| E_rsol | x3D | Solvation Correction Difference |
| E_rvdw | x3D | Van der Waals Interaction Energy |
| E_sol | i3D | Solvation energy |
| E_stb | i3D | Stretch-bend energy |
| E_str | i3D | Bond stretch energy |
| E_strain | i3D | E minus energy of local minimum |
| E_tor | i3D | Torsion energy |
| E_vdw | i3D | Van der Waals energy |
| FASA+ | i3D | Fractional positive accessible surface area |
| FASA- | i3D | Fractional negative accessible surface area |
| FASA_H | i3D | Fractional hydrophobic surface area |
| FASA_P | i3D | Fractional polar surface area |
| FCASA+ | i3D | Fractional charge-weighted positive surface area |
| FCASA- | i3D | Fractional charge-weighted negative surface area |
| FCharge | 2D | Sum of formal charges |
| GCUT_PEOE_0 | 2D | PEOE Charge GCUT (0/3) |
| GCUT_PEOE_1 | 2D | PEOE Charge GCUT (1/3) |
| GCUT_PEOE_2 | 2D | PEOE Charge GCUT (2/3) |
| GCUT_PEOE_3 | 2D | PEOE Charge GCUT (3/3) |
| GCUT_SLOGP_0 | 2D | LogP GCUT (0/3) |
| GCUT_SLOGP_1 | 2D | LogP GCUT (1/3) |
| GCUT_SLOGP_2 | 2D | LogP GCUT (2/3) |
| GCUT_SLOGP_3 | 2D | LogP GCUT (3/3) |
| GCUT_SMR_0 | 2D | Molar Refractivity GCUT (0/3) |
| GCUT_SMR_1 | 2D | Molar Refractivity GCUT (1/3) |
| GCUT_SMR_2 | 2D | Molar Refractivity GCUT (2/3) |
| GCUT_SMR_3 | 2D | Molar Refractivity GCUT (3/3) |
| glob | i3D | Molecular globularity |
| h_ema | 2D | Sum of EHT acceptor strengths |
| h_emd | 2D | Sum of EHT donor strengths |
| h_emd_C | 2D | Sum of EHT carbon donor strengths |
| h_logD | 2D | Octanol/water distribution coefficient (pH=7) |
| h_logP | 2D | Octanol/water partition coefficient |
| h_logS | 2D | Log solubility in water |
| h_log_dbo | 2D | Sum of log (1 + d-bond orders) |
| h_log_pbo | 2D | Sum of log (1 + p-bond orders) |
| h_mr | 2D | Molar Refractivity |
| h_pavgQ | 2D | Average total charge (pH=7) |
| h_pKa | 2D | Acidity (pH=7) |
| h_pKb | 2D | Basicity (pH=7) |
| h_pstates | 2D | Entropic state count (pH=7) |
| h_pstrain | 2D | Protonation state strain energy (pH=7) |
| Kier1 | 2D | First kappa shape index |
| Kier2 | 2D | Second kappa shape index |
| Kier3 | 2D | Third kappa shape index |
| KierA1 | 2D | First alpha modified shape index |
| KierA2 | 2D | Second alpha modified shape index |
| KierA3 | 2D | Third alpha modified shape index |
| KierFlex | 2D | Molecular flexibility |
| lip_acc | 2D | Lipinski Acceptor Count |
| lip_don | 2D | Lipinski Donor Count |
| lip_druglike | 2D | Lipinski Druglike Test |
| lip_violation | 2D | Lipinski Violation Count |
| logP(o/w) | 2D | Log octanol/water partition coefficient |
| logS | 2D | Log Solubility in Water |
| MNDO_dipole | i3D | Dipole moment |
| MNDO_E | i3D | Total energy (kcal/mol) |
| MNDO_Eele | i3D | Electronic energy (kcal/mol) |
| MNDO_HF | i3D | Heat of formation (kcal) |
| MNDO_HOMO | i3D | HOMO energy (eV) |
| MNDO_IP | i3D | Ionization potential (eV) |
| MNDO_LUMO | i3D | LUMO energy (eV) |
| mr | 2D | Molar refractivity |
| mutagenic | 2D | Mutagenicity |
| nmol | 2D | Number of molecules |
| npr1 | i3D | Normalized PMI ratio (1) (pmi1 / pmi3) |
| npr2 | i3D | Normalized PMI ratio (2) (pmi2 / pmi3) |
| opr_brigid | 2D | Oprea Rigid Bond Count |
| opr_leadlike | 2D | Oprea Leadlike Test |
| opr_nring | 2D | Oprea Ring Count |
| opr_nrot | 2D | Oprea Rotatable Bond Count |
| opr_violation | 2D | Oprea Violation Count |
| PC+ | 2D | Total positive partial charge |
| PC- | 2D | Total negative partial charge |
| PEOE_PC+ | 2D | Total positive partial charge |
| PEOE_PC- | 2D | Total negative partial charge |
| PEOE_RPC+ | 2D | Relative positive partial charge |
| PEOE_RPC- | 2D | Relative negative partial charge |
| PEOE_VSA+0 | 2D | Total positive 0 vdw surface area |
| PEOE_VSA+1 | 2D | Total positive 1 vdw surface area |
| PEOE_VSA+2 | 2D | Total positive 2 vdw surface area |
| PEOE_VSA+3 | 2D | Total positive 3 vdw surface area |
| PEOE_VSA+4 | 2D | Total positive 4 vdw surface area |
| PEOE_VSA+5 | 2D | Total positive 5 vdw surface area |
| PEOE_VSA+6 | 2D | Total positive 6 vdw surface area |
| PEOE_VSA-0 | 2D | Total negative 0 vdw surface area |
| PEOE_VSA-1 | 2D | Total negative 1 vdw surface area |
| PEOE_VSA-2 | 2D | Total negative 2 vdw surface area |
| PEOE_VSA-3 | 2D | Total negative 3 vdw surface area |
| PEOE_VSA-4 | 2D | Total negative 4 vdw surface area |
| PEOE_VSA-5 | 2D | Total negative 5 vdw surface area |
| PEOE_VSA-6 | 2D | Total negative 6 vdw surface area |
| PEOE_VSA_FHYD | 2D | Fractional hydrophobic vdw surface area |
| PEOE_VSA_FNEG | 2D | Fractional negative vdw surface area |
| PEOE_VSA_FPNEG | 2D | Fractional polar negative vdw surface area |
| PEOE_VSA_FPOL | 2D | Fractional polar vdw surface area |
| PEOE_VSA_FPOS | 2D | Fractional positive vdw surface area |
| PEOE_VSA_FPPOS | 2D | Fractional polar positive vdw surface area |
| PEOE_VSA_HYD | 2D | Total hydrophobic vdw surface area |
| PEOE_VSA_NEG | 2D | Total negative vdw surface area |
| PEOE_VSA_PNEG | 2D | Total polar negative vdw surface area |
| PEOE_VSA_POL | 2D | Total polar vdw surface area |
| PEOE_VSA_POS | 2D | Total positive vdw surface area |
| PEOE_VSA_PPOS | 2D | Total polar positive vdw surface area |
| petitjean | 2D | (diameter - radius) / diameter |
| petitjeanSC | 2D | (diameter - radius) / radius |
| PM3_dipole | i3D | Dipole moment |
| PM3_E | i3D | Total energy (kcal/mol) |
| PM3_Eele | i3D | Electronic energy (kcal/mol) |
| PM3_HF | i3D | Heat of formation (kcal) |
| PM3_HOMO | i3D | HOMO energy (eV) |
| PM3_IP | i3D | Ionization potential (eV) |
| PM3_LUMO | i3D | LUMO energy (eV) |
| pmi | i3D | Principal moment of inertia |
| pmi1 | i3D | Principal moment of inertia (1) |
| pmi2 | i3D | Principal moment of inertia (2) |
| pmi3 | i3D | Principal moment of inertia (3) |
| pmiX | x3D | Principal moment of inertia (X) |
| pmiY | x3D | Principal moment of inertia (Y) |
| pmiZ | x3D | Principal moment of inertia (Z) |
| pro_app_charge | Protein | Protein Charge at Debye Length |
| pro_asa_hph | Protein | Hydrophilic Surface Area |
| pro_asa_hyd | Protein | Hydrophobic Surface Area |
| pro_asa_vdw | Protein | Accessible Surface Area (Water Probe) |
| pro_coeff_280 | Protein | Extinction coefficient at 280 nm |
| pro_coeff_diff | Protein | Diffusion Coefficient |
| pro_coeff_fric | Protein | Frictional Coefficient |
| pro_debye | Protein | Debye Screening Length |
| pro_dipole_moment | Protein | Protein Dipole Moment |
| pro_eccen | Protein | Protein Eccentricity |
| pro_helicity | Protein | Protein Helix Ratio |
| pro_henry | Protein | Henry's Function f(ka) |
| pro_hyd_moment | Protein | Hydrophobicity Moment |
| pro_mass | Protein | Protein Mass in kDa |
| pro_mobility | Protein | Protein Mobility |
| pro_net_charge | Protein | Protein Net Charge |
| pro_patch_cdr_hyd | Protein | Area of hydrophobic protein patch(es) near CDRs |
| pro_patch_cdr_hyd_1 | Protein | Area of largest hydrophobic protein patch(es) near CDRs |
| pro_patch_cdr_hyd_2 | Protein | Area of 2 largest hydrophobic protein patch(es) near CDRs |
| pro_patch_cdr_hyd_3 | Protein | Area of 3 largest hydrophobic protein patch(es) near CDRs |
| pro_patch_cdr_hyd_4 | Protein | Area of 4 largest hydrophobic protein patch(es) near CDRs |
| pro_patch_cdr_hyd_5 | Protein | Area of 5 largest hydrophobic protein patch(es) near CDRs |
| pro_patch_cdr_hyd_n | Protein | Count of hydrophobic protein patch(es) near CDRs |
| pro_patch_cdr_ion | Protein | Area of ionic protein patch(es) near CDRs |
| pro_patch_cdr_ion_1 | Protein | Area of largest ionic protein patch(es) near CDRs |
| pro_patch_cdr_ion_2 | Protein | Area of 2 largest ionic protein patch(es) near CDRs |
| pro_patch_cdr_ion_3 | Protein | Area of 3 largest ionic protein patch(es) near CDRs |
| pro_patch_cdr_ion_4 | Protein | Area of 4 largest ionic protein patch(es) near CDRs |
| pro_patch_cdr_ion_5 | Protein | Area of 5 largest ionic protein patch(es) near CDRs |
| pro_patch_cdr_ion_n | Protein | Count of ionic protein patch(es) near CDRs |
| pro_patch_cdr_neg | Protein | Area of negative protein patch(es) near CDRs |
| pro_patch_cdr_neg_1 | Protein | Area of largest negative protein patch(es) near CDRs |
| pro_patch_cdr_neg_2 | Protein | Area of 2 largest negative protein patch(es) near CDRs |
| pro_patch_cdr_neg_3 | Protein | Area of 3 largest negative protein patch(es) near CDRs |
| pro_patch_cdr_neg_4 | Protein | Area of 4 largest negative protein patch(es) near CDRs |
| pro_patch_cdr_neg_5 | Protein | Area of 5 largest negative protein patch(es) near CDRs |
| pro_patch_cdr_neg_n | Protein | Count of negative protein patch(es) near CDRs |
| pro_patch_cdr_pos | Protein | Area of positive protein patch(es) near CDRs |
| pro_patch_cdr_pos_1 | Protein | Area of largest positive protein patch(es) near CDRs |
| pro_patch_cdr_pos_2 | Protein | Area of 2 largest positive protein patch(es) near CDRs |
| pro_patch_cdr_pos_3 | Protein | Area of 3 largest positive protein patch(es) near CDRs |
| pro_patch_cdr_pos_4 | Protein | Area of 4 largest positive protein patch(es) near CDRs |
| pro_patch_cdr_pos_5 | Protein | Area of 5 largest positive protein patch(es) near CDRs |
| pro_patch_cdr_pos_n | Protein | Count of positive protein patch(es) near CDRs |
| pro_patch_hyd | Protein | Area of hydrophobic protein patch(es) |
| pro_patch_hyd_1 | Protein | Area of largest hydrophobic protein patch(es) |
| pro_patch_hyd_2 | Protein | Area of 2 largest hydrophobic protein patch(es) |
| pro_patch_hyd_3 | Protein | Area of 3 largest hydrophobic protein patch(es) |
| pro_patch_hyd_4 | Protein | Area of 4 largest hydrophobic protein patch(es) |
| pro_patch_hyd_5 | Protein | Area of 5 largest hydrophobic protein patch(es) |
| pro_patch_hyd_n | Protein | Count of hydrophobic protein patch(es) |
| pro_patch_ion | Protein | Area of ionic protein patch(es) |
| pro_patch_ion_1 | Protein | Area of largest ionic protein patch(es) |
| pro_patch_ion_2 | Protein | Area of 2 largest ionic protein patch(es) |
| pro_patch_ion_3 | Protein | Area of 3 largest ionic protein patch(es) |
| pro_patch_ion_4 | Protein | Area of 4 largest ionic protein patch(es) |
| pro_patch_ion_5 | Protein | Area of 5 largest ionic protein patch(es) |
| pro_patch_ion_n | Protein | Count of ionic protein patch(es) |
| pro_patch_neg | Protein | Area of negative protein patch(es) |
| pro_patch_neg_1 | Protein | Area of largest negative protein patch(es) |
| pro_patch_neg_2 | Protein | Area of 2 largest negative protein patch(es) |
| pro_patch_neg_3 | Protein | Area of 3 largest negative protein patch(es) |
| pro_patch_neg_4 | Protein | Area of 4 largest negative protein patch(es) |
| pro_patch_neg_5 | Protein | Area of 5 largest negative protein patch(es) |
| pro_patch_neg_n | Protein | Count of negative protein patch(es) |
| pro_patch_pos | Protein | Area of positive protein patch(es) |
| pro_patch_pos_1 | Protein | Area of largest positive protein patch(es) |
| pro_patch_pos_2 | Protein | Area of 2 largest positive protein patch(es) |
| pro_patch_pos_3 | Protein | Area of 3 largest positive protein patch(es) |
| pro_patch_pos_4 | Protein | Area of 4 largest positive protein patch(es) |
| pro_patch_pos_5 | Protein | Area of 5 largest positive protein patch(es) |
| pro_patch_pos_n | Protein | Count of positive protein patch(es) |
| pro_pI_3D | Protein | Structure-based pI Prediction |
| pro_pI_seq | Protein | Sequence-based pI Prediction |
| pro_r_gyr | Protein | Radius of Gyration |
| pro_r_solv | Protein | Hydrodynamic Radius |
| pro_sed_const | Protein | Sedimentation Constant |
| pro_volume | Protein | Protein Volume |
| pro_zdipole | Protein | Zeta Dipole Moment |
| pro_zeta | Protein | Zeta potential at Debye Length |
| pro_zquadrupole | Protein | Zeta Quadrupole Moment |
| Q_PC+ | 2D | Total positive partial charge |
| Q_PC- | 2D | Total negative partial charge |
| Q_RPC+ | 2D | Relative positive partial charge |
| Q_RPC- | 2D | Relative negative partial charge |
| Q_VSA_FHYD | 2D | Fractional hydrophobic vdw surface area |
| Q_VSA_FNEG | 2D | Fractional negative vdw surface area |
| Q_VSA_FPNEG | 2D | Fractional polar negative vdw surface area |
| Q_VSA_FPOL | 2D | Fractional polar vdw surface area |
| Q_VSA_FPOS | 2D | Fractional positive vdw surface area |
| Q_VSA_FPPOS | 2D | Fractional polar positive vdw surface area |
| Q_VSA_HYD | 2D | Total hydrophobic vdw surface area |
| Q_VSA_NEG | 2D | Total negative vdw surface area |
| Q_VSA_PNEG | 2D | Total polar negative vdw surface area |
| Q_VSA_POL | 2D | Total polar vdw surface area |
| Q_VSA_POS | 2D | Total positive vdw surface area |
| Q_VSA_PPOS | 2D | Total polar positive vdw surface area |
| radius | 2D | Smallest vertex eccentricity in graph |
| reactive | 2D | Reactivity |
| rgyr | i3D | Radius of gyration |
| rings | 2D | Number of rings |
| RPC+ | 2D | Relative positive partial charge |
| RPC- | 2D | Relative negative partial charge |
| rsynth | 2D | Synthetic Feasibility |
| SlogP | 2D | Log Octanol/Water Partition Coefficient |
| SlogP_VSA0 | 2D | Bin 0 SlogP (-10 ,-0.40] |
| SlogP_VSA1 | 2D | Bin 1 SlogP (-0.40,-0.20] |
| SlogP_VSA2 | 2D | Bin 2 SlogP (-0.20, 0.00] |
| SlogP_VSA3 | 2D | Bin 3 SlogP ( 0.00, 0.10] |
| SlogP_VSA4 | 2D | Bin 4 SlogP ( 0.10, 0.15] |
| SlogP_VSA5 | 2D | Bin 5 SlogP ( 0.15, 0.20] |
| SlogP_VSA6 | 2D | Bin 6 SlogP ( 0.20, 0.25] |
| SlogP_VSA7 | 2D | Bin 7 SlogP ( 0.25, 0.30] |
| SlogP_VSA8 | 2D | Bin 8 SlogP ( 0.30, 0.40] |
| SlogP_VSA9 | 2D | Bin 9 SlogP ( 0.40,10] |
| SMR | 2D | Molar Refractivity |
| SMR_VSA0 | 2D | Bin 0 SMR (0.000,0.110] |
| SMR_VSA1 | 2D | Bin 1 SMR (0.110,0.260] |
| SMR_VSA2 | 2D | Bin 2 SMR (0.260,0.350] |
| SMR_VSA3 | 2D | Bin 3 SMR (0.350,0.390] |
| SMR_VSA4 | 2D | Bin 4 SMR (0.390,0.440] |
| SMR_VSA5 | 2D | Bin 5 SMR (0.440,0.485] |
| SMR_VSA6 | 2D | Bin 6 SMR (0.485,0.560] |
| SMR_VSA7 | 2D | Bin 7 SMR (0.560,10] |
| std_dim1 | i3D | Standard dimension 1 |
| std_dim2 | i3D | Standard dimension 2 |
| std_dim3 | i3D | Standard dimension 3 |
| TPSA | 2D | Topological Polar Surface Area (A**2) |
| VAdjEq | 2D | Vertex adjacency information (equal) |
| VAdjMa | 2D | Vertex adjacency information (mag) |
| VDistEq | 2D | Vertex distance equality index |
| VDistMa | 2D | Vertex distance magnitude index |
| vdw_area | 2D | Van der Waals surface area (A**2) |
| vdw_vol | 2D | Van der Waals volume (A**3) |
| vol | i3D | Van der Waals volume |
| VSA | i3D | Van der Waals surface area |
| vsa_acc | 2D | VDW acceptor surface area (A**2) |
| vsa_acid | 2D | VDW acidic surface area (A**2) |
| vsa_base | 2D | VDW basic surface area (A**2) |
| vsa_don | 2D | VDW donor surface area (A**2) |
| vsa_hyd | 2D | VDW hydrophobe surface area (A**2) |
| vsa_other | 2D | VDW other surface area (A**2) |
| vsa_pol | 2D | VDW polar surface area (A**2) |
| vsurf_A | i3D | Amphiphilic moment |
| vsurf_CP | i3D | Critical packing parameter |
| vsurf_CW1 | i3D | Capacity factor at -0.2 |
| vsurf_CW2 | i3D | Capacity factor at -0.5 |
| vsurf_CW3 | i3D | Capacity factor at -1.0 |
| vsurf_CW4 | i3D | Capacity factor at -2.0 |
| vsurf_CW5 | i3D | Capacity factor at -3.0 |
| vsurf_CW6 | i3D | Capacity factor at -4.0 |
| vsurf_CW7 | i3D | Capacity factor at -5.0 |
| vsurf_CW8 | i3D | Capacity factor at -6.0 |
| vsurf_D1 | i3D | Hydrophobic volume at -0.2 |
| vsurf_D2 | i3D | Hydrophobic volume at -0.4 |
| vsurf_D3 | i3D | Hydrophobic volume at -0.6 |
| vsurf_D4 | i3D | Hydrophobic volume at -0.8 |
| vsurf_D5 | i3D | Hydrophobic volume at -1.0 |
| vsurf_D6 | i3D | Hydrophobic volume at -1.2 |
| vsurf_D7 | i3D | Hydrophobic volume at -1.4 |
| vsurf_D8 | i3D | Hydrophobic volume at -1.6 |
| vsurf_DD12 | i3D | vsurf_EDmin1, vsurf_EDmin2 distance |
| vsurf_DD13 | i3D | vsurf_EDmin1, vsurf_EDmin3 distance |
| vsurf_DD23 | i3D | vsurf_EDmin2, vsurf_EDmin3 distance |
| vsurf_DW12 | i3D | vsurf_EWmin1, vsurf_EWmin2 distance |
| vsurf_DW13 | i3D | vsurf_EWmin1, vsurf_EWmin3 distance |
| vsurf_DW23 | i3D | vsurf_EWmin2, vsurf_EWmin3 distance |
| vsurf_EDmin1 | i3D | Lowest hydrophobic energy |
| vsurf_EDmin2 | i3D | 2nd lowest hydrophobic energy |
| vsurf_EDmin3 | i3D | 3rd lowest hydrophobic energy |
| vsurf_EWmin1 | i3D | Lowest hydrophilic energy |
| vsurf_EWmin2 | i3D | 2nd lowest hydrophilic energy |
| vsurf_EWmin3 | i3D | 3rd lowest hydrophilic energy |
| vsurf_G | i3D | Surface globularity |
| vsurf_HB1 | i3D | H-bond donor capacity at -0.2 |
| vsurf_HB2 | i3D | H-bond donor capacity at -0.5 |
| vsurf_HB3 | i3D | H-bond donor capacity at -1.0 |
| vsurf_HB4 | i3D | H-bond donor capacity at -2.0 |
| vsurf_HB5 | i3D | H-bond donor capacity at -3.0 |
| vsurf_HB6 | i3D | H-bond donor capacity at -4.0 |
| vsurf_HB7 | i3D | H-bond donor capacity at -5.0 |
| vsurf_HB8 | i3D | H-bond donor capacity at -6.0 |
| vsurf_HL1 | i3D | First hydrophilic-lipophilic balance |
| vsurf_HL2 | i3D | Second hydrophilic-lipophilic balance |
| vsurf_ID1 | i3D | Hydrophobic integy moment at -0.2 |
| vsurf_ID2 | i3D | Hydrophobic integy moment at -0.4 |
| vsurf_ID3 | i3D | Hydrophobic integy moment at -0.6 |
| vsurf_ID4 | i3D | Hydrophobic integy moment at -0.8 |
| vsurf_ID5 | i3D | Hydrophobic integy moment at -1.0 |
| vsurf_ID6 | i3D | Hydrophobic integy moment at -1.2 |
| vsurf_ID7 | i3D | Hydrophobic integy moment at -1.4 |
| vsurf_ID8 | i3D | Hydrophobic integy moment at -1.6 |
| vsurf_IW1 | i3D | Hydrophilic integy moment at -0.2 |
| vsurf_IW2 | i3D | Hydrophilic integy moment at -0.5 |
| vsurf_IW3 | i3D | Hydrophilic integy moment at -1.0 |
| vsurf_IW4 | i3D | Hydrophilic integy moment at -2.0 |
| vsurf_IW5 | i3D | Hydrophilic integy moment at -3.0 |
| vsurf_IW6 | i3D | Hydrophilic integy moment at -4.0 |
| vsurf_IW7 | i3D | Hydrophilic integy moment at -5.0 |
| vsurf_IW8 | i3D | Hydrophilic integy moment at -6.0 |
| vsurf_R | i3D | Surface rugosity |
| vsurf_S | i3D | Interaction field area |
| vsurf_V | i3D | Interaction field volume |
| vsurf_W1 | i3D | Hydrophilic volume at -0.2 |
| vsurf_W2 | i3D | Hydrophilic volume at -0.5 |
| vsurf_W3 | i3D | Hydrophilic volume at -1.0 |
| vsurf_W4 | i3D | Hydrophilic volume at -2.0 |
| vsurf_W5 | i3D | Hydrophilic volume at -3.0 |
| vsurf_W6 | i3D | Hydrophilic volume at -4.0 |
| vsurf_W7 | i3D | Hydrophilic volume at -5.0 |
| vsurf_W8 | i3D | Hydrophilic volume at -6.0 |
| vsurf_Wp1 | i3D | Polar volume at -0.2 |
| vsurf_Wp2 | i3D | Polar volume at -0.5 |
| vsurf_Wp3 | i3D | Polar volume at -1.0 |
| vsurf_Wp4 | i3D | Polar volume at -2.0 |
| vsurf_Wp5 | i3D | Polar volume at -3.0 |
| vsurf_Wp6 | i3D | Polar volume at -4.0 |
| vsurf_Wp7 | i3D | Polar volume at -5.0 |
| vsurf_Wp8 | i3D | Polar volume at -6.0 |
| Weight | 2D | Molecular weight (CRC) |
| weinerPath | 2D | Weiner path number |
| weinerPol | 2D | Weiner polarity number |
| zagreb | 2D | Zagreb index |

### Performance: AR

| **Supplementary Table 2: DECTRE. Run-time: 3 minutes 33 seconds. Performance: 92.73%.** | | | | |
| --- | --- | --- | --- | --- |
|  | True Agonist | True Antag. | True Decoy | Precision |
| Pred. Agonist | 1415 | 239 | 0 | 85.55% |
| Pred. Antag. | 85 | 313 | 0 | 78.64% |
| Pred. Decoy | 79 | 301 | 7254 | 95.02% |
| Recall | 89.61% | 39.69% | 100.00% |  |

| **Supplementary Table 3: NAIBAY. Run-time: 1 minute 9 seconds. Performance: 94.64%.** | | | | |
| --- | --- | --- | --- | --- |
|  | True Agonist | True Antag. | True Decoy | Precision |
| Pred. Agonist | 1434 | 243 | 2 | 85.41% |
| Pred. Antag. | 67 | 482 | 1 | 87.64% |
| Pred. Decoy | 78 | 128 | 7251 | 97.24% |
| Recall | 90.82% | 56.51% | 99.96% |  |

| **Supplementary Table 4: NEUNET. Run-time: 6 hours 34 minutes 37 seconds. Performance: 94.09%.** | | | | |
| --- | --- | --- | --- | --- |
|  | True Agonist | True Antag. | True Decoy | Precision |
| Pred. Agonist | 1492 | 218 | 0 | 87.25% |
| Pred. Antag. | 84 | 370 | 2 | 81.14% |
| Pred. Decoy | 3 | 265 | 7252 | 96.44% |
| Recall | 94.49% | 43.38% | 99.97% |  |

| **Supplementary Table 5: RANFOR. Run-time: 2 hours 21 minutes 18 seconds. Performance: 91.41%.** | | | | |
| --- | --- | --- | --- | --- |
|  | True Agonist | True Antag. | True Decoy | Precision |
| Pred. Agonist | 1232 | 184 | 2 | 86.88% |
| Pred. Antag. | 347 | 370 | 0 | 51.60% |
| Pred. Decoy | 0 | 299 | 7252 | 96.04% |
| Recall | 78.02% | 43.38% | 99.97% |  |

| **Supplementary Table 6: SVM. Run-time: 1 day 16 hours 47 minutes 47 seconds. Performance: 77.83%.** | | | | |
| --- | --- | --- | --- | --- |
|  | True Agonist | True Antag. | True Decoy | Precision |
| Pred. Agonist | 286 | 0 | 1 | 99.65% |
| Pred. Antag. | 0 | 0 | 0 | 00.00% |
| Pred. Decoy | 1293 | 853 | 7253 | 77.17% |
| Recall | 18.11% | 00.00% | 99.99% |  |

### Performance: ER

| **Supplementary Table 7: DECTRE. Run-time: 13 minutes 39 seconds. Performance: 92.61%.** | | | | |
| --- | --- | --- | --- | --- |
|  | True Agonist | True Antag. | True Decoy | Precision |
| Pred. Agonist | 1283 | 405 | 1 | 75.96% |
| Pred. Antag. | 122 | 504 | 4 | 80.00% |
| Pred. Decoy | 12 | 428 | 10400 | 95.94% |
| Recall | 90.54% | 37.70% | 99.95% |  |

| **Supplementary Table 8: NAIBAY. Run-time: 2 minutes 25 seconds. Performance: 90.51%.** | | | | |
| --- | --- | --- | --- | --- |
|  | True Agonist | True Antag. | True Decoy | Precision |
| Pred. Agonist | 336 | 55 | 0 | 85.93% |
| Pred. Antag. | 1081 | 1176 | 7 | 51.94% |
| Pred. Decoy | 0 | 106 | 10398 | 98.99% |
| Recall | 23.71% | 87.96% | 99.93% |  |

| **Supplementary Table 9: NEUNET. Run-time: 21 hours 38 minutes 12 seconds. Performance: 95.59%.** | | | | |
| --- | --- | --- | --- | --- |
|  | True Agonist | True Antag. | True Decoy | Precision |
| Pred. Agonist | 965 | 40 | 0 | 96.02% |
| Pred. Antag. | 452 | 1213 | 4 | 72.68% |
| Pred. Decoy | 0 | 84 | 10401 | 99.20% |
| Recall | 68.10% | 90.73% | 99.96% |  |

| **Supplementary Table 10: RANFOR. Run-time: 8 hours 1 minute 19 seconds. Performance: 83.43%.** | | | | |
| --- | --- | --- | --- | --- |
|  | True Agonist | True Antag. | True Decoy | Precision |
| Pred. Agonist | 171 | 36 | 0 | 82.61% |
| Pred. Antag. | 1198 | 406 | 4 | 25.25% |
| Pred. Decoy | 48 | 895 | 10401 | 91.69% |
| Recall | 12.07% | 30.37% | 99.96% |  |

| **Supplementary Table 11: SVM. Run-time: 15 days 12 hours 40 minutes 35 seconds. Performance: 81.39%.** | | | | |
| --- | --- | --- | --- | --- |
|  | True Agonist | True Antag. | True Decoy | Precision |
| Pred. Agonist | 58 | 5 | 0 | 92.06% |
| Pred. Antag. | 2 | 250 | 3 | 98.04% |
| Pred. Decoy | 1357 | 1082 | 10402 | 81.01% |
| Recall | 04.09% | 18.70% | 99.97% |  |

### Performance: GR

| **Supplementary Table 12: DECTRE. Run-time: 1 minute 35 seconds. Performance: 96.08%.** | | | | |
| --- | --- | --- | --- | --- |
|  | True Agonist | True Antag. | True Decoy | Precision |
| Pred. Agonist | 3437 | 16 | 0 | 99.54% |
| Pred. Antag. | 0 | 26 | 0 | 100.00% |
| Pred. Decoy | 0 | 435 | 7594 | 94.58% |
| Recall | 100.00% | 05.45% | 100.00% |  |

| **Supplementary Table 13: NAIBAY. Run-time: 1 minute 6 seconds. Performance: 96.02%.** | | | | |
| --- | --- | --- | --- | --- |
|  | True Agonist | True Antag. | True Decoy | Precision |
| Pred. Agonist | 3437 | 283 | 1 | 92.37% |
| Pred. Antag. | 0 | 26 | 6 | 81.25% |
| Pred. Decoy | 0 | 168 | 7587 | 97.83% |
| Recall | 100.00% | 05.45% | 99.91% |  |

| **Supplementary Table 14: NEUNET. Run-time: 8 hours 46 minutes 1 second. Performance: 96.32%.** | | | | |
| --- | --- | --- | --- | --- |
|  | True Agonist | True Antag. | True Decoy | Precision |
| Pred. Agonist | 3437 | 0 | 6 | 99.83% |
| Pred. Antag. | 0 | 59 | 0 | 100.00% |
| Pred. Decoy | 0 | 418 | 7588 | 94.78% |
| Recall | 100.00% | 12.37% | 99.92% |  |

| **Supplementary Table 15: RANFOR. Run-time: 4 hours 20 minutes 41 seconds. Performance: 96.03%.** | | | | |
| --- | --- | --- | --- | --- |
|  | True Agonist | True Antag. | True Decoy | Precision |
| Pred. Agonist | 3437 | 16 | 6 | 99.36% |
| Pred. Antag. | 0 | 26 | 0 | 100.00% |
| Pred. Decoy | 0 | 435 | 7588 | 94.58% |
| Recall | 100.00% | 05.45% | 99.92% |  |

| **Supplementary Table 16: SVM. Run-time: 13 days 22 hours 40 minutes 27 seconds. Performance: 80.93%.** | | | | |
| --- | --- | --- | --- | --- |
|  | True Agonist | True Antag. | True Decoy | Precision |
| Pred. Agonist | 252 | 0 | 6 | 97.67% |
| Pred. Antag. | 0 | 0 | 1 | 00.00% |
| Pred. Decoy | 1165 | 1337 | 10398 | 80.60% |
| Recall | 17.78% | 00.00% | 99.93% |  |

### Performance: PR

| **Supplementary Table 17: DECTRE. Run-time: 2 minutes 58 seconds. Performance: 99.92%.** | | | | |
| --- | --- | --- | --- | --- |
|  | True Agonist | True Antag. | True Decoy | Precision |
| Pred. Agonist | 793 | 0 | 0 | 100.00% |
| Pred. Antag. | 6 | 258 | 1 | 97.36% |
| Pred. Decoy | 0 | 0 | 7908 | 100.00% |
| Recall | 99.25% | 100.00% | 99.99% |  |

| **Supplementary Table 18: NAIBAY. Run-time: 1 minute. Performance: 99.91%.** | | | | |
| --- | --- | --- | --- | --- |
|  | True Agonist | True Antag. | True Decoy | Precision |
| Pred. Agonist | 795 | 0 | 3 | 99.62% |
| Pred. Antag. | 0 | 258 | 1 | 99.61% |
| Pred. Decoy | 4 | 0 | 7905 | 99.95% |
| Recall | 99.50% | 100.00% | 99.95% |  |

| **Supplementary Table 19: NEUNET. Run-time: 4 hours 14 minutes 59 seconds. Performance: 98.64%.** | | | | |
| --- | --- | --- | --- | --- |
|  | True Agonist | True Antag. | True Decoy | Precision |
| Pred. Agonist | 789 | 112 | 0 | 87.57% |
| Pred. Antag. | 6 | 146 | 0 | 96.05% |
| Pred. Decoy | 4 | 0 | 7909 | 99.95% |
| Recall | 98.75% | 56.59% | 100.00% |  |

| **Supplementary Table 20: RANFOR. Run-time: 1 hour 19 minutes 57 seconds. Performance: 99.87%.** | | | | |
| --- | --- | --- | --- | --- |
|  | True Agonist | True Antag. | True Decoy | Precision |
| Pred. Agonist | 789 | 0 | 1 | 99.87% |
| Pred. Antag. | 6 | 258 | 1 | 97.36% |
| Pred. Decoy | 4 | 0 | 7907 | 99.95% |
| Recall | 98.75% | 100.00% | 99.97% |  |

| **Supplementary Table 21: SVM. Run-time: 6 days 2 hours 55 minutes 33 seconds. Performance: 91.36%.** | | | | |
| --- | --- | --- | --- | --- |
|  | True Agonist | True Antag. | True Decoy | Precision |
| Pred. Agonist | 184 | 0 | 0 | 100.00% |
| Pred. Antag. | 46 | 100 | 2 | 67.57% |
| Pred. Decoy | 569 | 158 | 7907 | 91.58% |
| Recall | 23.03% | 38.76% | 99.97% |  |
